# Timelapse and volumetric imaging of mitochondrial networking using NAD(P)H autofluorescence via 2-photon microscopy

**DOI:** 10.1101/2025.01.15.633075

**Authors:** Blanche ter Hofstede, Alex J. Walsh

**Author notes:** Address all correspondence to Alex J. Walsh.

## Abstract

**Significance:** Mitochondria are dynamic organelles that play a key role in energy production and maintaining cellular homeostasis. The regulation of mitochondrial dynamics, involving both fission and fusion, is vital for maintaining a healthy population of mitochondria within the cell. Alterations in mitochondrial dynamics have been associated with various disease states, such as metabolic and neurodegenerative diseases and cancer.

**Aim:** We describe a protocol for imaging and analyzing NAD(P)H intensity to visualize the movement of mitochondria over time and in 3D to visualize the distribution of the mitochondrial network within the cell.

**Approach:** A multiphoton (MP) laser scanning microscope was used to image NAD(P)H autofluorescence signal of MDA-MB-231 cells at 750 nm excitation. A mitochondrial fluorescent dye, MitoSpy Orange, was used to validate the signal. Laser power, image size, dwell time, interval time, and imaging duration were optimized for timelapse and 3D imaging to minimize photodamage and maximize autofluorescence signal.

**Results:** The NAD(P)H signal in 2D was imaged with a frame rate of 0.4 frames per second (FPS) allowing for visualization of mitochondria movement. 3D imaging was performed with a frame rate of 0.5 FPS for a single cell with a thickness of approximately 15 microns that allowed for visualization of the mitochondria network within the cell.

**Conclusions:** This protocol motivates using label-free imaging techniques to study mitochondrial dynamics in a non-destructive manner, suitable for drug screening and understanding the effects of mitochondrial dynamic alterations in disease.

## 1 Introduction

Mitochondria are dynamic organelles responsible for ATP production via oxidative phosphorylation (OXPHOS). They play a critical role in cell homeostasis, maintenance, and cell death ^1,2^. To maintain a healthy mitochondria pool and provide energy to the cell, mitochondria are highly motive organelles that undergo fission and fusion as well as biogenesis and mitophagy^3^. This process is referred to as mitochondrial dynamics. Typically, mitochondrial fusion leads to networking of mitochondria and is associated with increased OXPHOS, while mitochondrial fission is associated with fragmented mitochondria, proliferation, mitophagy, and, when in excess, increased ROS production^4^. Mitochondrial dynamics and its dysfunction have been heavily studied in applications such as metabolic and neurodegenerative diseases and cancer^5,6^

However, analysis of mitochondria dynamics is limited to end-point assays that quantify fission and fusion proteins or destructive imaging techniques that require fluorescent dyes. Both time-lapse and 3D imaging have been achieved with confocal imaging, although limited due to high photobleaching and phototoxicity potential from out-of-focus laser light. Multiphoton laser scanning microscopy allows for reduced photodamage and photobleaching due to the focusing of excitation light and lower energy wavelengths used. Autofluorescence from NAD(P)H has been shown to correlate with mitochondrial signal due to the high concentration of NAD(P)H within mitochondria and the long lifetime of unquenched protein-bound state of NAD(P)H that occurs in the mitochondria. Mitochondria have been imaged using autofluorescence at different scales ranging from the macroscopic *in vivo*^7^ to quantifying individual mitochondrial movement *in vitro* using autofluorescence intensity and fluorescence lifetime microscopy (FLIM)^8,9^. Yet, a gap remains for imaging volumetric spatial information or temporal information about the mitochondrial movement at the cellular level using autofluorescence intensity.

In this paper, we describe a protocol for imaging and analyzing NAD(P)H intensity to track and quantify mitochondrial networks over time to visualize the movement of mitochondria and in 3D to visualize the distribution of the mitochondrial networking in the cell. The mitochondria signal of NAD(P)H in MDA-MB-231 triple-negative breast cancer cells is validated using a mitochondrial stain MitoSpy Orange. Timelapse and 3D images were taken using a laser scanning multiphoton microscope imaged at a frame rate of 0.4 FPS to achieve resolution and speeds to track the mitochondria while minimizing photobleaching and photodamage. Label-free analysis using autofluorescence signal of mitochondria could allow for a better understanding of mitochondrial dynamics in disease and could be used for drug screening to see the effects of drugs on mitochondrial networking and morphology.

## 2 Materials and Methods

### 2.1 Cell Culture

MDA-MB-231 cells were maintained in 10 mL of RPMI 1640 (Gibco, 11879020) supplemented with 10% fetal bovine serum, 1% antibiotic-antimycotic, and 5 mM glucose in T-75 flasks. Cells were incubated at 37°C with 5% CO_2_ and 95% humidified air. Cell media was exchanged every 2 days after rinsing with PBS. Once reaching 70-80% confluency (∼ 1 week), cells were passaged.

### 2.2 Sample preparation

Once MDA-MB-231 cells reached 70-80% confluency, cells were seeded on either 35 mm glass bottom imaging dish (MatTek) or a glass bottom 6 well plate (MatTek) at a density of approximately 10^5^ cells in cell growth media. The samples were incubated at 37°C with 5% CO_2_ and 95% humidified air for 24-48 hours until reaching a confluency of 70-80%. Prior to imaging, cells were rinsed with PBS, and warm cell growth media was added to the dish and incubated for 30 minutes.

### 2.3 Instrumentation and processing for visualization of mitochondria NAD(P) autofluorescence

All images were taken on a custom-built laser-scanning multiphoton microscope (Marianas, 3i) with a tunable Ti: sapphire femtosecond laser (Coherent, Chameleon Ultra II). NAD(P)H images were captured using an excitation wavelength of 750 nm on a photomultiplier (PMT) (Hamamatsu) with bandpass filters of 447/60 nm and 550/88 nm. Images were taken with either a 100X 1.4 NA (Zeiss) or a 150X 1.34 NA glycerine objectives (Zeiss). Samples were imaged in a stage-top incubator (OkoLabs) at 37°C with 5% CO_2_ and 95% humidified air. A power meter (Thorlabs) was used in the light path with a 30% split to ensure laser power was consistent with autofluorescence intensity images across acquisitions. The power measurements described here are the power recorded at this location. This reading was mapped with power meter values at the imaging field of view for both objectives to ensure the power did not reach values greater than 5 mW to prevent photodamage to the sample. SlideBook 24 (3i) was used as the image acquisition software. Images were pre-processed in FIJI, where after deconvolution with a simulated point spread function (PSF), a Gaussian blur filter and despeckling filters were applied to the timelapse and 3D images.

### 2.4 Mitochondrial signal validation

For verification of NADH and mitochondria signal, a fluorescent mitochondrial dye was used. mitochondrial staining, cells were left to incubate for 30 minutes in 100 nM MitoSpy Orange (BioLegend) before imaging. Cells were rinsed with warm PBS twice and warm cell growth media was added.

Using an excitation of 750 nm, NAD(P)H and MitoSpy signals were imaged on two separate PMTs spectrally separated using appropriate emission filters. The gain for the PMTs were set to 85% and 65% for the NAD(P)H and MitoSpy signals, respectively, due to the higher quantum yield of the fluorescent dye. Using an image size of 1024×1024, images were taken at an excitation power of less than 25 mW.

### 2.5 Timelapse Imaging of Mitochondrial Dynamics

The following protocol has been optimized to capture the movement of the mitochondria of MDA-MB-231 cells in 2D while maximizing NAD(P)H autofluorescent signal and image resolution while minimizing photodamage. The susceptibility to photodamage and speed of mitochondria movement is cell type dependent, and timing should be tested and adjusted accordingly.

Timelapse images were obtained using either the 100X objective (0.10 µm/pixel) or 150X (0.067 µm/pixel) objectives. A pixel dwell time of 2 us, an image resolution of 1024×1024, and a detector gain of 85% were used to resolve the mitochondria. The resulting frame rate was 0.4 frames per second (FPS) if the full scanning field was used. NAD(P)H autofluorescence was excited at 750 nm with a power of approximately 15 mW for time intervals of less than 1 second and a power of less than 20 mW for intervals between 1-2 seconds.

### 2.6 3D Visualization of Mitochondria Networks Using Autofluorescence

For 3D imaging, the power could be increased to achieve higher signal. Using the 100X or 150X objective, a single cell or clump of cells was imaged by cropping the scanning field to increase imaging speeds. The pixel dwell time was set to 2 µs. The Z-stack step size was approximately 0.2 microns to satisfy the Nyquist frequency and maximize imaging speeds to reduce mitochondria blurring. The resulting frame rate was approximately 0.5 FPS for a 1024×1024 image size. For single cells and clumps of cells, the scan field was reduced to increase the frame rate (Fig. 1). For the 100X objective, a single cell was approximately 93.5 × 67.9 µm^2^ in X-Y for a frame rate of 0.5 FPS. The NAD(P)H signal was excited at 750 nm with an average power of approximately 20 mW.

**Figure 1.**
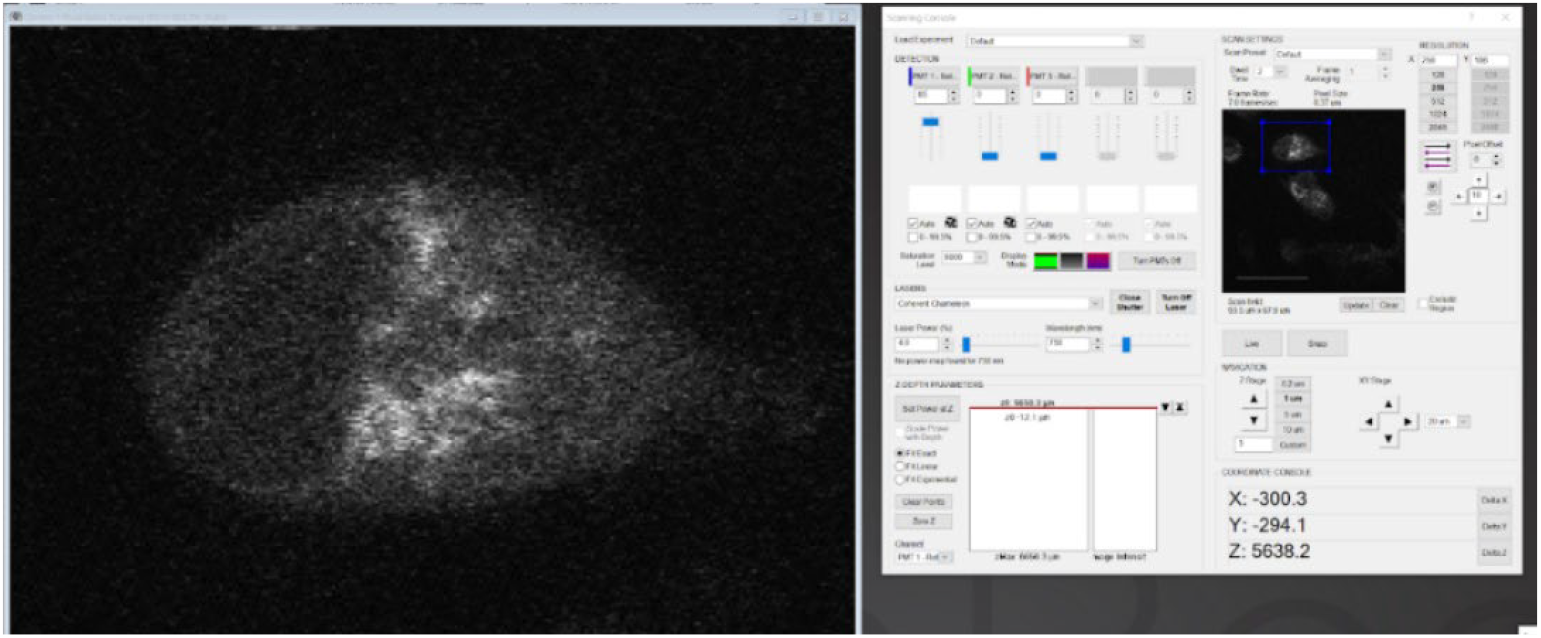
Live view image (256×256 image size for focusing on cell) of single MDA-MB-231 cell cropped in Slidebook 24 for 3D autofluorescence imaging.

## 3 Results

### 3.1 Mitochondrial labeling validation using mitochondrial fluorescent dye

To show the ability to image mitochondria using NAD(P)H autofluorescence, a mitochondrial stain was used, and the two signals were imaged simultaneously. MDA-MB-231 cells were labeled with MitoSpy Orange allowing for spectral separation using 2 emission filters. The signal overlap (Fig. 2) shows the higher NAD(P)H autofluorescence intensity pixels corresponding to the MitoSpy Orange signal. As expected, due to the lower quantum yield of NAD(P)H, the signal of NAD(P)H is lower than the MitoSpy dye even with the higher PMT detector gain. Discrepancies in signal could be due to dye distribution caused by low membrane mitochondria potential, a lack of NAD(P)H in the mitochondria, or other sources of autofluorescence, such as lipofuscin.

**Figure 2.**
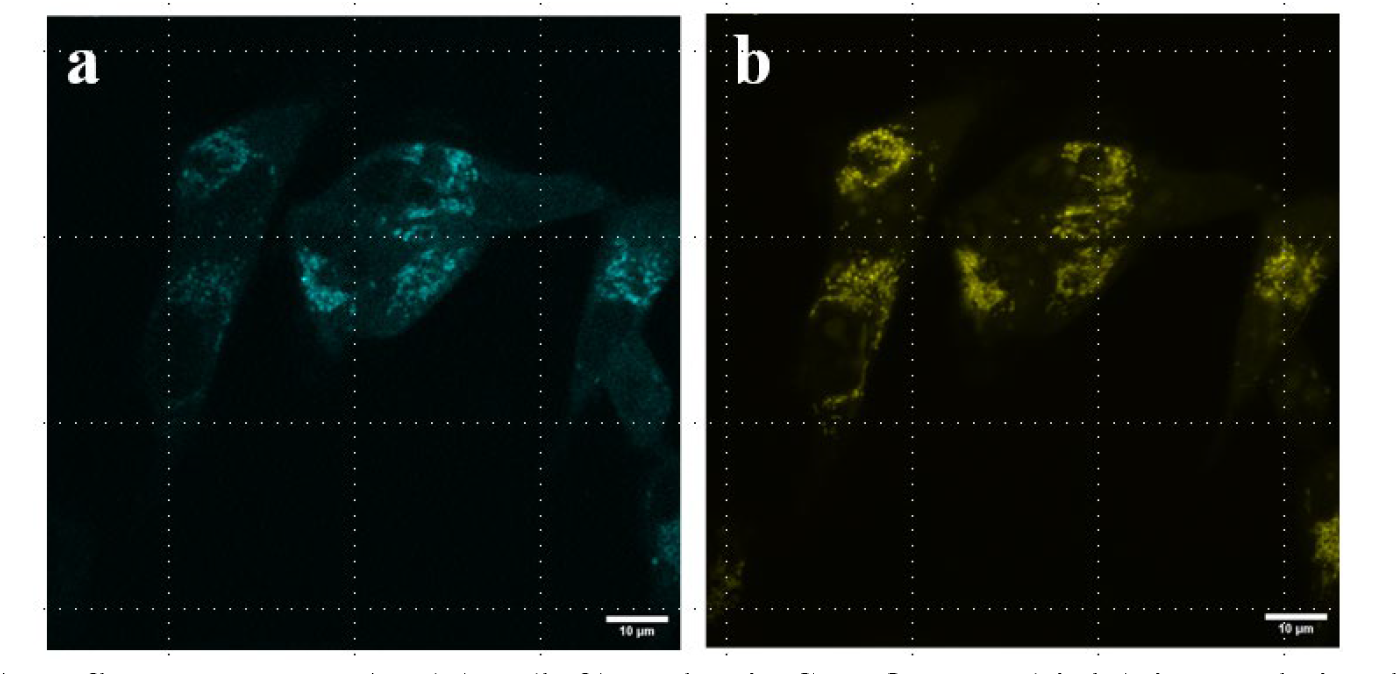
Autofluorescence NAD(P)H (left) and MitoSpy Orange (right) imaged simultaneously at 750 nm with PMT gains of 85% and 60%, respectively. Scale bar = 10 µm.

### 3.2 Timelapse visualization of mitochondria in 2D

Timelapse imaging of autofluorescence from NAD(P)H allowed for visualization of mitochondria movement at imaging speeds of 0.4 FPS with high image resolution as shown in Fig. 3. A montage of a cropped section (Fig. 3b) of the image demonstrated where the movement of the NAD(P)H corresponding to mitochondria is tracked over 50 frames with a frame rate of 0.4 FPS. This allows for visualizing the movement for quantification using tracking algorithms.

**Figure 3.**
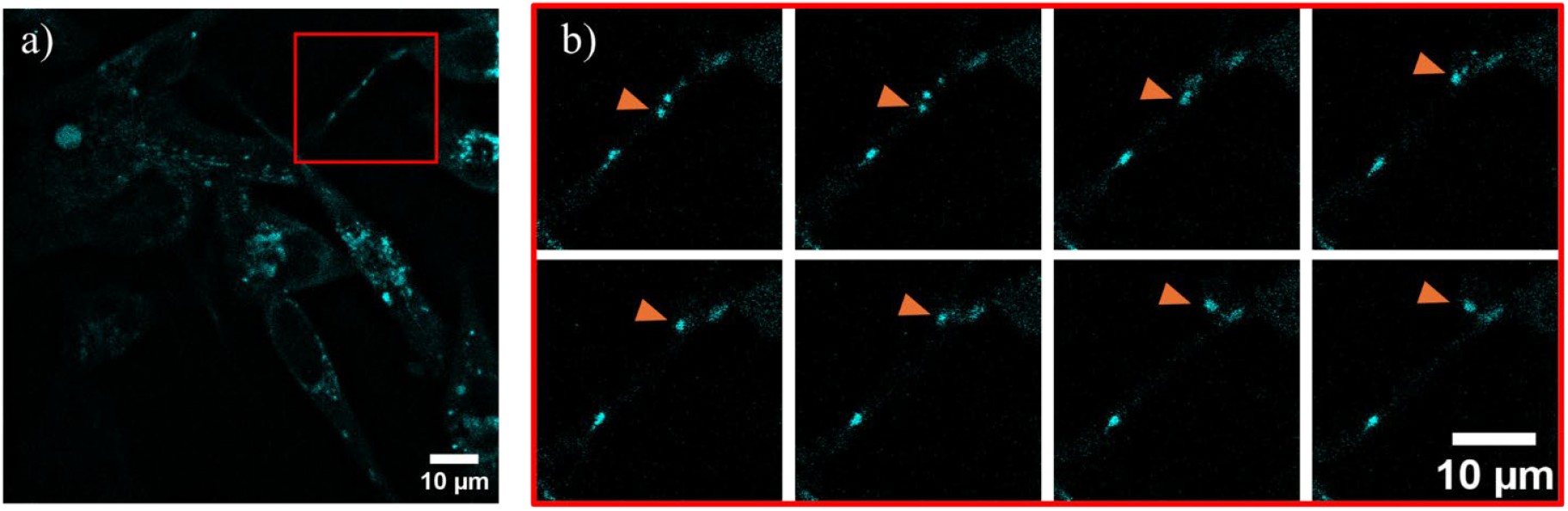
Representative timelapse images of NAD(P)H signal with a time interval of 100 ms at 0.4 FPS. (a) Full field of view of cluster of cells. (b) Timelapse montage cropped view to visualize mitochondria movement with every 4 frames shown.

### 3.3 3D visualization of mitochondria network in single cells

3D imaging using mitochondrial autofluorescence allowed for better visualization of the mitochondrial networking within the cell. Fig. 4A shows a maximum intensity project of a single cell imaged at a frame rate of 0.5 FPS. Compared to the single XY cross section in Fig 4B from the same Z-stack, more information is gathered about the distribution of the mitochondria in the cell.

**Figure 4.**
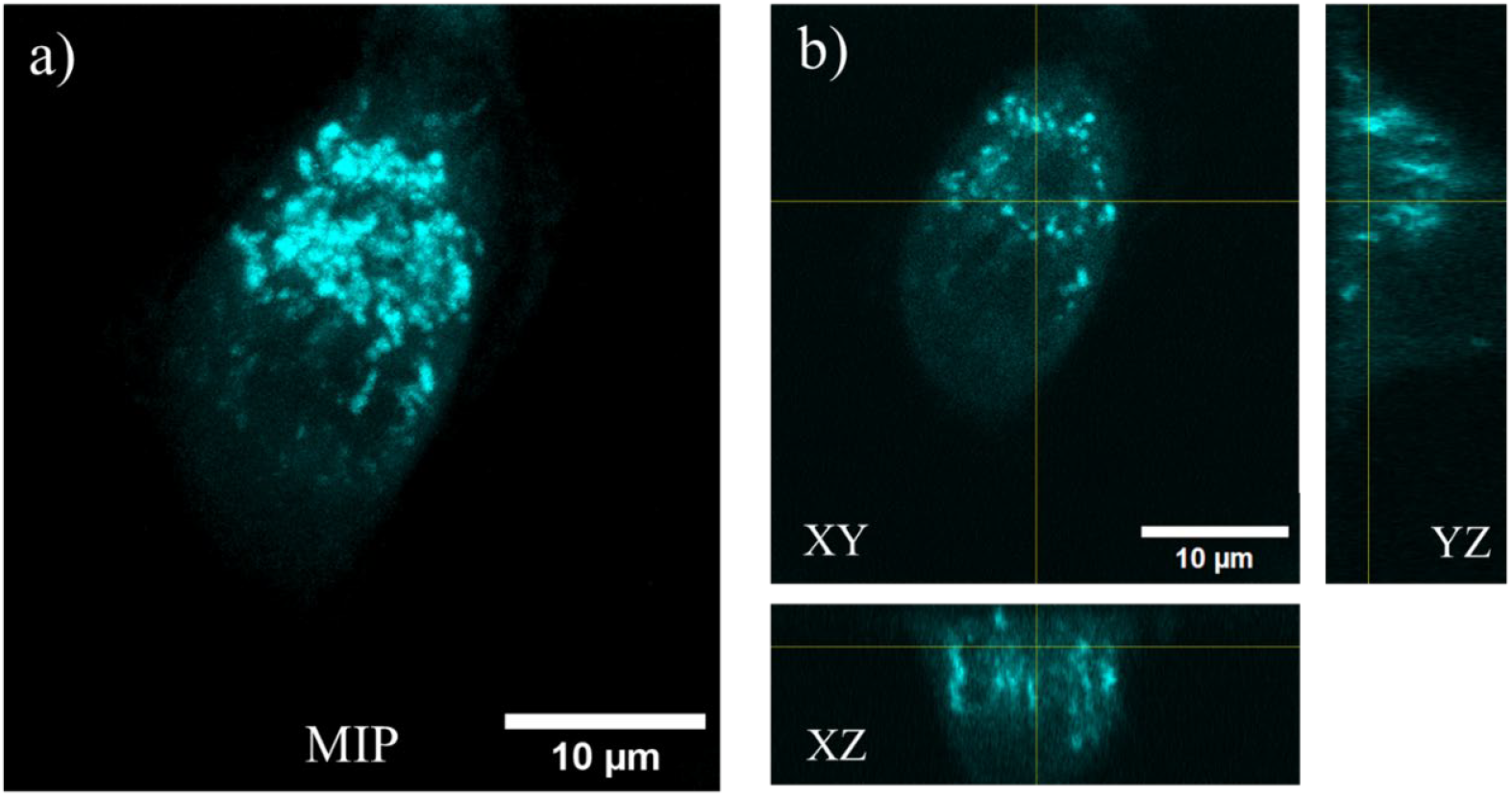
Representative 3D image of MDA-MB-231 cells. (A) Maximum intensity project of image stack. (B) Orthogonal views of single cell. Imaged at 0.5 FPS with voxel size of 0.038×0.038×0.24 µm^3^.

## 4 Discussion

We have outlined a protocol for imaging mitochondrial NAD(P)H autofluorescence signal. The high-intensity NAD(P)H signal was correlated with a mitochondrial fluorescent dye MitoSpy as shown in Fig. 2. Areas of mismatch with the MitoSpy signal were observed that could be signal contributed by lipofuscin that exhibit autofluorescence that spectrally overlaps with NAD(P)H^10^. Timelapse and 3D images were taken with parameters optimized for faster imaging while still capturing the mitochondria networking and minimizing photodamage. Volumetric imaging allowed for better visualization, although it is limited by the speed of the laser scanning microscope. Minimizing the scanning field helped increase the speeds but some dynamics may be missed. Mitochondria vary in size, shape, speed, and degree of networking depending on the cell type and state of the cell^11^. This protocol could be applied to other cell types and applications. Laser power, PMT gain, and imaging time interval may need to be adjusted for other cell types. The next step in this work is to test the protocol on several different cells, such as muscle cells, neurons, and MCF10A, an epithelial cell line typically used as a control for MDA-MB-231 cells. Overall, this protocol motivates using label-free imaging techniques to study mitochondrial dynamics in a non-destructive manner, suitable for drug screening and understanding the effects of mitochondrial dynamic alterations in disease.

## Disclosures

The authors declare no conflicts of interest.

## Acknowledgments

Funding sources include Chan Zuckerberg Initiative (2022-251380), NIH NIGMS (R35 GM142990), and Texas A&M University. We thank Gloria Echeverria and Steven W. Wall for providing the MDA-MB-231 cells.

## Notes

### Competing Interest Statement

The authors have declared no competing interest.

